# Prevalent Presence of Periodic Actin-spectrin-based Membrane Skeleton in a Broad Range of Neuronal Cell Types and Animal Species

**DOI:** 10.1101/045856

**Authors:** Jiang He, Ruobo Zhou, Zhuhao Wu, Monica Carrasco, Peri Kurshan, Jonathan Farley, David Simon, Guiping Wang, Boran Han, Junjie Hao, Evan Heller, Marc Freeman, Kang Shen, Tom Maniatis, Marc Tessier-Lavigne, Xiaowei Zhuang

**Author notes:** Current Address: Institute of Medical Engineering and Sciences, Massachusetts Institute of Technology, Cambridge, MA, 02139, USA. Current Address: Department of Biomedical Science, Faculty of Health Sciences, University of Talca, Talca, 3460000, Chile. These authors contributed equally to this work.

## Abstract

Actin, spectrin and associated molecules form a periodic, sub-membrane cytoskeleton in the axons of neurons. For a better understanding of this membrane-associated periodic skeleton (MPS), it is important to address how prevalent this structure is in different neuronal types, different subcellular compartments, and across different animal species. Here, we investigated the organization of spectrin in a variety of neuronal and glial-cell types. We observed the presence of MPS in all of the tested neuronal types cultured from mouse central and peripheral nervous systems, including excitatory and inhibitory neurons from several brain regions, as well as sensory and motor neurons. Quantitative analyses show that MPS is preferentially formed in axons in all neuronal types tested here: spectrin shows a long-range, periodic distribution throughout all axons, but only appears periodic in a small fraction of dendrites, typically in the form of isolated patches in sub-regions of these dendrites. As in dendrites, we also observed patches of periodic spectrin structures in a small fraction of glial-cell processes in four types of glial cells cultured from rodent tissues. Interestingly, despite its strong presence in the axonal shaft, MPS is absent in most presynaptic boutons, but is present in a substantial fraction of dendritic spine necks, including some projecting from dendrites where such a periodic structure is not observed in the shaft. Finally, we found that spectrin is capable of adopting a similar periodic organization in neurons of a variety of animal species, including *Caenorhabditis elegans*, *Drosophila*, *Gallus gallus*, *Mus musculus* and *Homo sapiens*.

## Introduction

Actin is critically involved in the regulation of neuronal polarization, differentiation and growth of neuronal processes, cargo trafficking, and plasticity of synapses (1–3). Spectrin is an actin-binding protein that is important for the development and stabilization of axons, and maintenance of neuronal polarization (4–6). In *C. elegans*, spectrin is important for the stability and integrity of axons under mechanical stress (4, 6) and for mechanosensation (6), and spectrin depletion results in axon breakage during animal locomotion (4). In *Drosophila*, spectrin has been shown to be involved in axonal path finding (7) and stabilization of presynaptic terminals (8). In mice, spectrin null mutations are embryonically lethal and neurons with spectrin knockdown display defects in axonal initial segment assembly (5, 9, 10).

Actin and spectrin form a two-dimensional polygonal lattice structure underneath the membrane of erythrocytes (11). Recently, we discovered a novel form of actin-spectrin-based sub-membrane skeleton structure in neuronal axons (12) using super-resolution STORM imaging (13, 14). This membrane-associated periodic skeleton (MPS) has been observed in both fixed and live cultured neurons (12, 15, 16) and in brain tissue sections (12). In this structure, short actin filaments are organized into repetitive, ring-like structures that wrap around the circumference of the axon with a periodicity of ~190 nm; adjacent actin rings are connected by spectrin tetramers and actin short filaments in the rings are capped by adducin(12). This structure also appears to organize other associated molecules, such as ankyrin and sodium channels, into a periodic distribution in axons (12, 16). During neuronal development, MPS originates from the axonal region proximal to the soma and propagates to distal axonal terminals (16). At a relatively late stage during development, specific isoforms of ankyrin and spectrin molecules, ankyrin-G and βIV spectrin, are recruited to the axon initial segment (AIS) (17, 18), and these molecules are also assembled into the MPS structure, adopting a similar periodic distribution (16, 19). As in the AIS, this periodic structure is also present in the nodes of Ranvier (20). This periodic skeletal structure has been shown to preferentially form in axons compared to dendrites in primary neuronal cultures: actin and spectrin typically form a long-range, periodic lattice structure throughout the entire axonal shaft, except for the very distal region near the growth cone, in essentially all observed axons. In contrast, such a periodic structure was observed in only a small fraction (~10-30%) of dendrites, and typically appeared as short, isolated patches in portions of these dendrites (16, 20). The local concentration of spectrin is a key determinant for the preferential formation of MPS in axons: in wild type neurons, βII spectrin is enriched in axons, and artificially increasing the concentration of βII spectrin through overexpression is sufficient to induce the formation of MPS in all dendrites (16). Ankyrin-B appears to be an important regulator of this structure: in ankyrin-B knockout mice, βII spectrin becomes evenly distributed between axons and dendrites, leading to the formation of the long-range MPS structure in all dendrites (16) without perturbing the MPS structure in axons (16, 21).

The ubiquitous expression of spectrin in the nervous systems of nearly all animal species (22) raises the question of how widespread the MPS structure is in different nervous system cell types and distinct subcellular compartments, and how conserved this structure is across different animal species. A recent paper reports the presence of the MPS structure in several nervous system cell types from rodents (23). Here we investigated these questions regarding the prevalence and conservation of the MPS structure by examining the distribution of spectrin in many different types of rodent neurons and glial cells, and across a variety of organisms ranging from *C. elegans* to *Homo sapiens.* Furthermore, we examined the distribution of spectrin in presynaptic and postsynaptic compartments of axons and dendrites, respectively, to shed light on the relation between the MPS structure and synapses.

## Results

### The MPS structure is present in both excitatory and inhibitory neurons cultured from mouse cortex, hippocampus and midbrain

We initially observed the membrane-associated, periodic actin-spectrin structure in axons of rodent hippocampal and cortical neurons (12). The neurons observed in this earlier study are most likely excitatory pyramidal neurons. To test whether our findings extend to other excitatory neuronal types, or inhibitory neurons in the mouse brain, we first cultured neurons from three different mouse brain regions, the cortex, hippocampus and midbrain, and distinguished excitatory and inhibitory neurons using immunofluorescence against vGlut1 and GAD2, respectively (Fig. S1). We then labeled βII spectrin in these neurons using immunofluorescence and imaged immunolabeled βII spectrin using 3D STORM imaging, a super-resolution imaging method that uses stochastic switching and high-precision localization of single molecules to obtain sub-diffraction-limit image resolution (13, 14). We distinguished axons from dendrites either by morphology, with the axon being a distinctly long process projecting from the soma, or by immunofluorescence, with dendrites being MAP2 positive. We observed highly periodic distributions of βII spectrin in the axons of nearly all (>90%) vGlut1+ excitatory neurons cultured from all three brain regions (Fig. 1A and Figs. S2A, S3A, S4A). Autocorrelation analysis indicated that the spacing was around 190 nm (Fig. 1C and Figs. S2A, S3A, S4A), consistent with the value determined previously (12, 16). βII spectrin also adopted a periodic distribution in the axons of nearly all (90%) GAD2+ inhibitory neurons from mouse cortex, hippocampus and midbrain (Fig. 1B and Figs. S2B, S3B, S4B). We employed autocorrelation analysis to systematically quantify the degree of periodicity for βII spectrin in axons of these neurons. The amplitude of the averaged autocorrelation function of many randomly selected segments of axons provided a measure of the degree of periodicity. As shown in Fig. 1C, both the spacing and the degree of periodicity in GAD2+ inhibitory neurons were quantitatively similar to those obtained for vGlut1+ excitatory neurons.

**Fig. 1.**
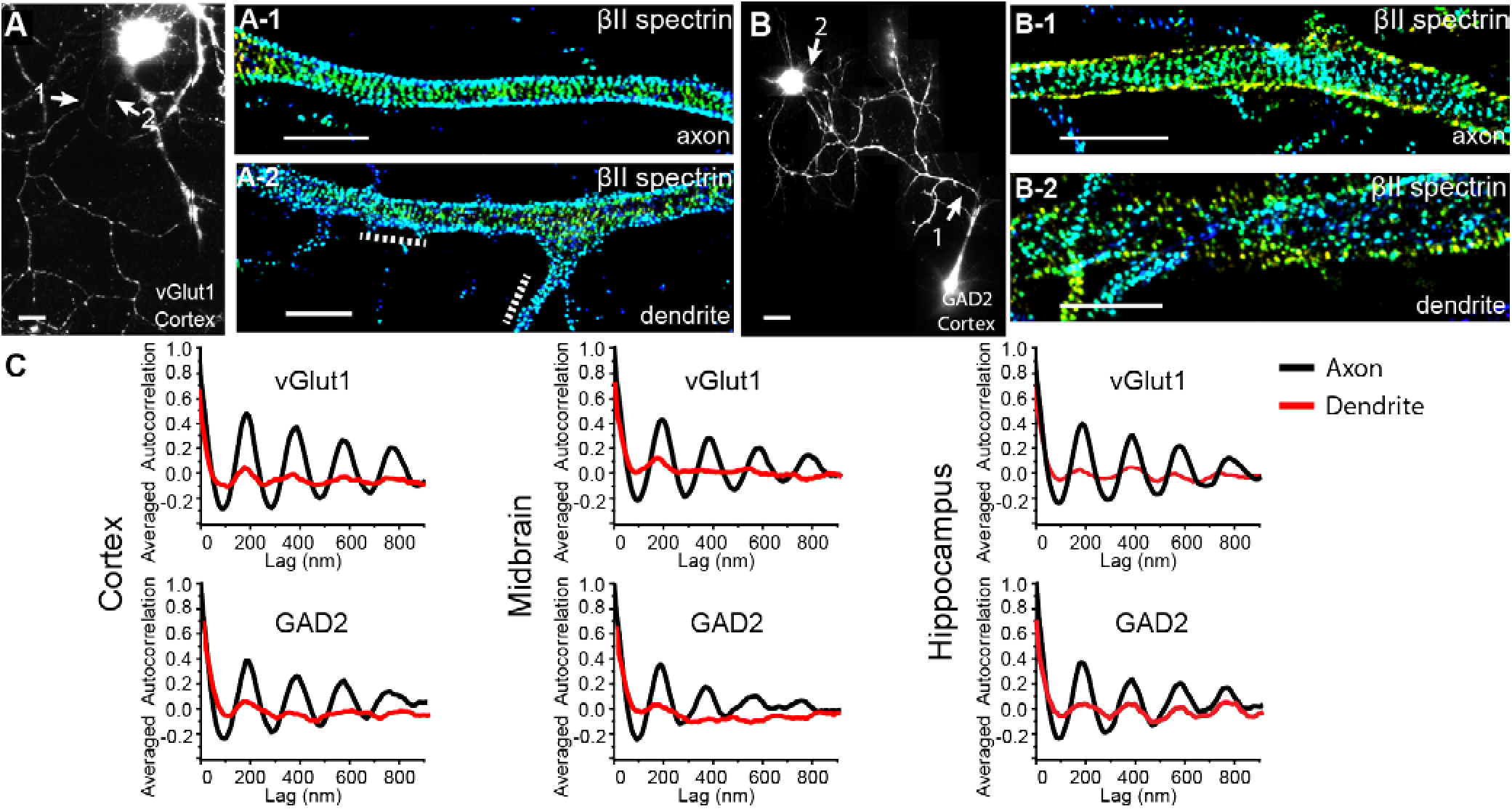
The MPS structure is present in both excitatory and inhibitory neurons from mouse cortex, midbrain and hippocampus. Cultured neurons were immunostained with vGlut1 or GAD2 to label the excitatory and inhibitory neurons, respectively. (A) Reconstructed image of a vGlut1+ cortical neuron (left) with 3D STORM images of βII spectrin in an axonal (A-1) and dendritic (A-2) region shown on the right. The axonal (A-1) and dendritic (A-2) regions correspond to the regions indicated by arrows in the left image. Dotted lines in (A-2) indicate patches of periodic pattern in the dendritic shaft. In 3D STORM images, localizations at different z values are depicted in different colors. (B) Similar to (A) but for a GAD2+ cortical neuron. (C) Averaged autocorrelation functions calculated from multiple, randomly selected axonal (black) and dendritic (red) regions in neurons cultured from cortex (N = 18 for axons and N = 9 for dendrites of GAD2+ neurons; N = 9 for axons and N = 8 for dendrites of vGlut1+ neurons), midbrain (N = 8 for axons and N = 7 for dendrites of GAD2+ neurons; N = 13 for axons and N = 11 for dendrites of vGlut1+ neurons) and hippocampus (N = 6 for axons and N = 8 for dendrites of GAD2+ neurons; N = 8 for axons and N = 9 for dendrites of vGlut1+ neurons), respectively. Scale bar: 20 μm for conventional images, 2 μm for STORM images.

In contrast, βII spectrin exhibited a much less regular distribution in dendritic shafts from the same neurons: unlike in axons, where βII spectrin adopted a long-range, periodic pattern throughout nearly the entire axonal shaft, the βII spectrin distribution exhibited a periodic pattern only occasionally in a small fraction of dendritic shafts, often as short, isolated patches (Fig. 1A–2 and Figs. S2A-2, S4A-2, S4B-2). This observation is similar to what we observed previously in mouse hippocampal neurons (16). The average amplitudes of the autocorrelation functions derived from many randomly selected segments of dendrites were much smaller than those observed for axons in both excitatory and inhibitory neurons from all three brain regions (Fig. 1C), quantitatively showing that the degree of periodicity in dendrites was much less than that in axons.

**Fig. 2.**
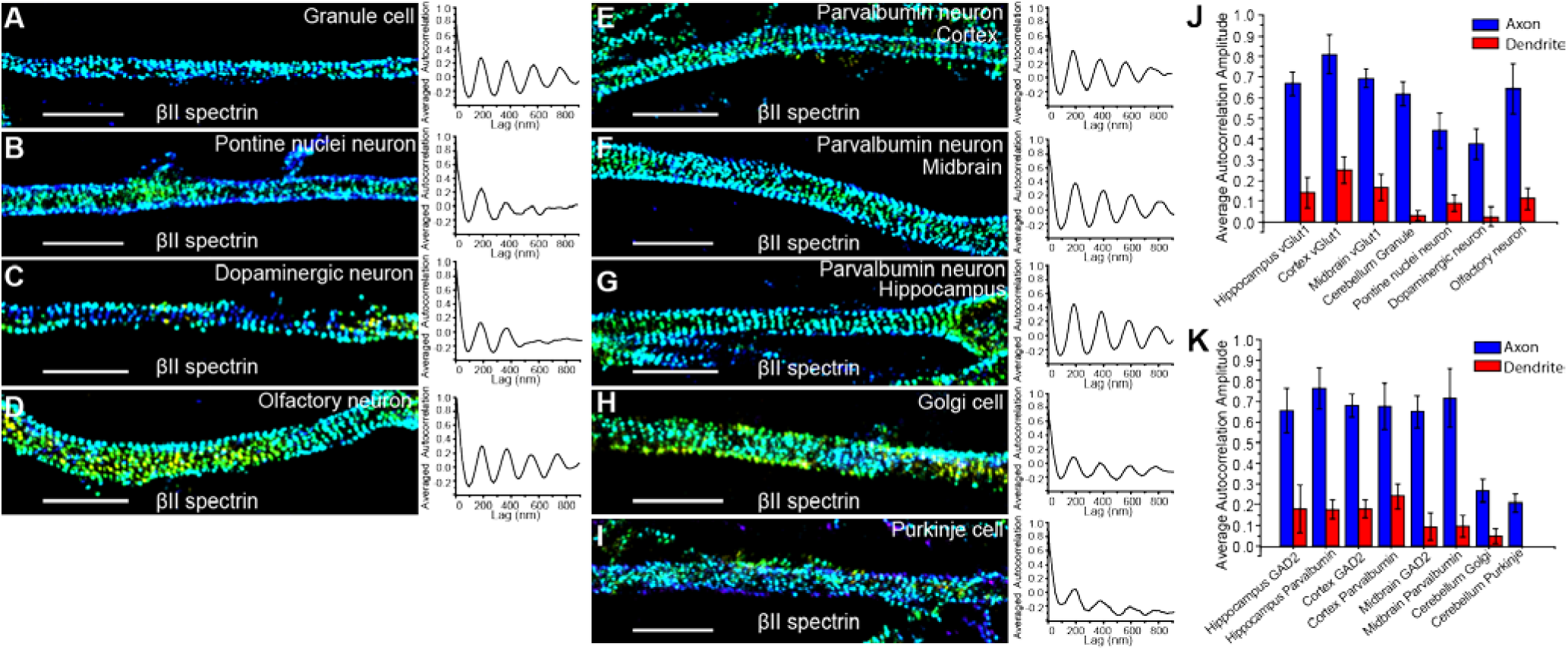
MPS is present in multiple subtypes of inhibitory and excitatory neurons in the mouse central nervous system. (A-I) Representative STORM images of βII spectrin for a selected axonal region (left) and averaged autocorrelation functions from multiple axons (right) for cultured cerebellar granule cells (A; N = 7 for averaged autocorrelation calculation), Pontine nuclei neurons (B; N = 7), Dopaminergic neurons (C; N = 9), Olfactory neurons (D; N = 16), Parvalbumin neurons from cortex (E; N = 9), Parvalbumin neurons from midbrain (F; N = 6), Parvalbumin neurons from hippocampus (G; N = 13), Golgi cells (H; N = 8) and Purkinje cells (I; N = 7) from cerebellum. (J and K) Averaged autocorrelation amplitudes of axons (blue) versus dendrites (red) for the neuronal subtypes we imaged in the mouse central nervous system. Error bars are SEM. Scale bar: 2μm.

### The MPS structure is present in multiple specific subtypes of excitatory and inhibitory neurons in the mouse central nervous system

The proteins vGlut1 and GAD2 are general markers for glutamatergic excitatory and GABAergic inhibitory neurons, respectively. Next, we selected some morphologically and functionally distinct subtypes of excitatory and inhibitory neurons from the mouse brain for further investigation. Specifically, for excitatory neurons, we imaged cerebellar granule cells, dopaminergic neurons from the midbrain, and pontine nuclei excitatory neurons. For inhibitory neurons, we imaged Golgi cells and Purkinje cells from cerebellum, as well as Parvalbumin positive cells from the cortex, midbrain and hippocampus. Granule cells and Purkinje cells were chosen because of their distinct morphologies. Granule cells are among the smallest cells in the brain, and Purkinje cells are unique in their large size and elaborate dendritic branches (24). Dopaminergic neurons release the neurotransmitter dopamine, and represent a type of excitatory neuron distinct from glutamatergic neurons(24). Parvalbumin cells belong to a specific subtype of GABAergic inhibitory neurons (24). In addition, we imaged olfactory neurons from the olfactory bulb.

Cultured dopaminergic neurons, olfactory neurons, Golgi cells, Purkinje cells and Parvalbumin cells were identified using immunostaining against tyrosine hydroxylase (25), olfactory marker protein (26), metabotropic glutamate receptors (27), calbindin (28) and parvalbumin, respectively (Fig. S1 and S5), while granule cells cultured from cerebellum were distinguished by their lack of calbindin staining and relatively small soma size. We also used general neuronal markers MAP2 and NeuN to exclude glial cells from our analysis. All neuronal subtypes were cultured for at least 10 days *in vitro* until a distinctly long axon had projected from the soma to ensure sufficient axon specification. The neurons were then fixed and immunostained with βII spectrin for STORM imaging. Axons were again distinguished from dendrites either by their morphology or lack of MAP2 staining. As shown in Fig. 2A-I, all of these neuronal subtypes exhibited a periodic distribution of βII spectrin in their axons, with a spacing of ~190 nm. Autocorrelation analyses showed that the degree of periodicity (i.e. autocorrelation amplitude) was similar among most of these excitatory and inhibitory neurons (Fig. 2J, K). Two exceptions were Golgi and Purkinje cells, which showed less regularity (Fig. 2H, I, K). This is probably due to a relatively slow development process for these cultured cerebellar neurons as evidenced by less elaborate dendrites observed for these neurons *in vitro* (29, 30). In addition, all of these specific neuronal subtypes showed a substantially higher propensity for MPS formation in axons with only occasional presence of periodic patterns in dendrites; concordantly, quantitative autocorrelation analyses showed substantially lower average autocorrelation amplitudes in dendrites (Fig. 2J, K). Together, our data suggest that the MPS structure is prevalent in morphologically and functionally diverse excitatory and inhibitory neuronal types.

As many neurons are myelinated in the brain, we next examined whether myelination had any effect on the formation of MPS using a previously developed neuron-glia co-culture system (31). Myelin sheaths were identified using an antibody against myelin basic protein (MBP). As shown by STORM images and autocorrelation analysis, βII spectrin was periodically distributed in both MBP positive and MBP negative axonal segments with a nearly identical degree of periodicity (Fig. S6), suggesting that myelination has little impact on the formation of the periodic membrane skeleton *in vitro*, consistent with the observation of the MPS structure in myelinated sciatic nerves (23).

### The MPS structure is present in mouse peripheral sensory and motor axons

The aforementioned results were all from neurons restricted to the mouse central nervous system. We next examined whether MPS was also present in peripheral neurons by studying two representative types, motor neurons and dorsal root ganglia (DRG) sensory neurons. Motor neurons were derived from mouse embryonic stem (mES) cells and differentiated motor neurons were identified by Hb9 staining (32). βII spectrin was again detected by immunostaining and STORM imaging. The distribution of βII spectrin was highly periodic in the axons, but mostly non-periodic in the dendrites of mES-derived motor neurons (Fig. 3A, B). Different from motor neurons, DRG sensory neurons are pseudounipolar neurons that have axons but lack dendrites (24). We imaged immunostained βII spectrin in explants of DRG sensory neurons (Fig. 3C) and in dissociated DRG sensory neurons (Fig. 3D); the periodic distribution of βII spectrin was clearly evident in the axons of these neurons as well.

**Fig. 3.**
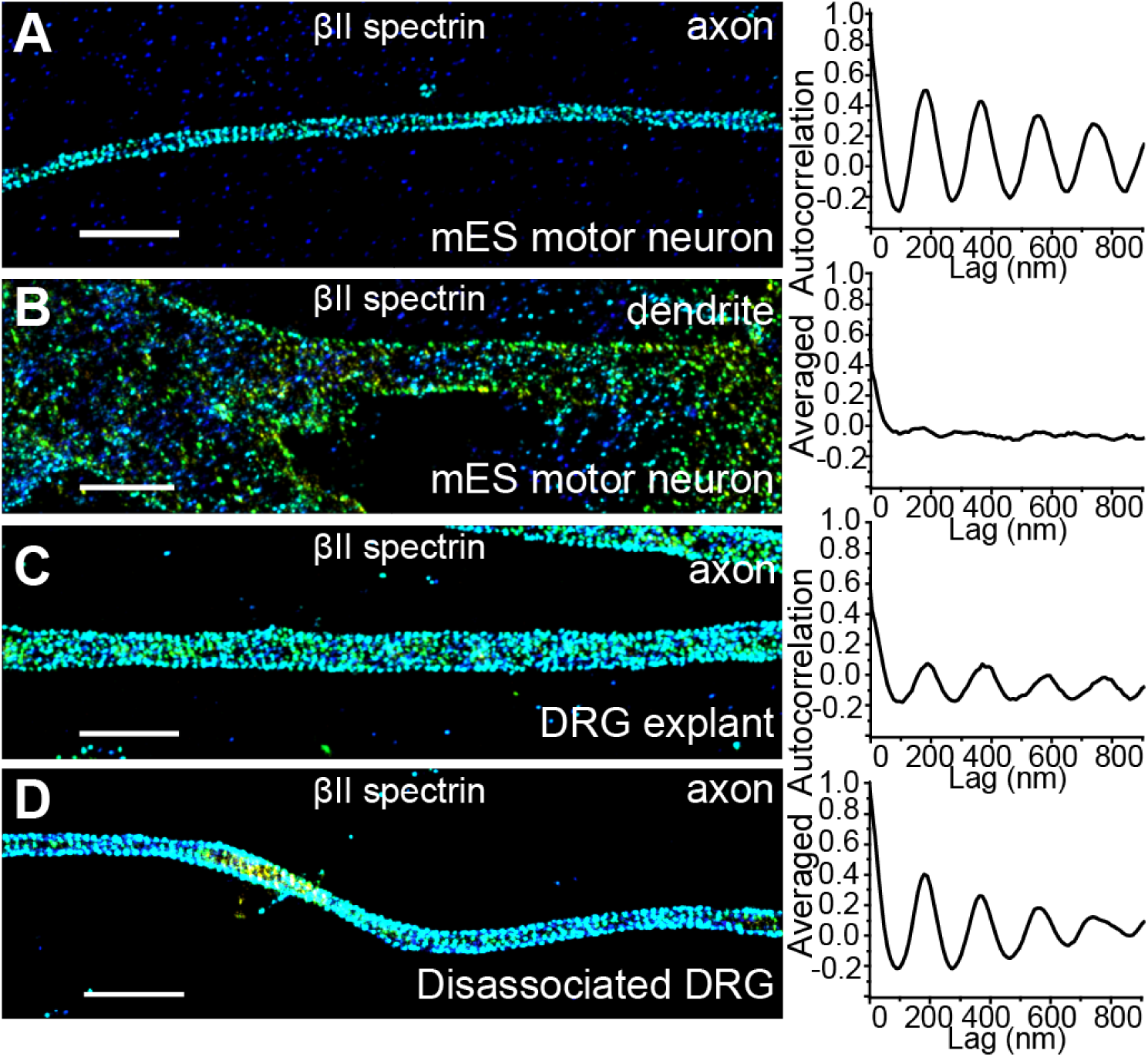
MPS is present in mouse peripheral sensory and motor axons. (A) Representative STORM images of βII spectrin for a selected axonal region (left) and averaged autocorrelation functions from multiple axons (right, N = 12) for mouse embryonic stem cell (mES)-derived motor neurons. (B) Same as in (A) but for dendrites in mES-derived motor neurons (N = 11). (C, D) Same as (A) but for axons in DRG neurons ((C) for explants, N = 13; (D) for dissociated DRG culture, N = 10). Scale bar: 2 μm.

### The distribution of βII spectrin in presynaptic boutons and dendritic spine necks

We next extended our investigation to two neuronal compartments in axons and dendrites related to synapses: axonal boutons (presynaptic) and dendritic spines (postsynaptic). Spectrin is expressed in synapses and plays important roles in synapse stabilization and spine morphogenesis (8, 33, 34). To image βII spectrin in presynaptic boutons, we transiently expressed a GFP-tagged αII spectrin (or GFP-tagged all spectrin) construct (16) in a sparse subpopulation of cultured hippocampal neurons, immununolabeled GFP-tagged spectrin with anti-GFP, and performed STORM imaging at DIV 14. The presynaptic boutons were marked by immunofluorescence of Bassoon and imaged at the diffraction-limited resolution. Such transient expression was performed to allow for the spectrin labeling in only a sparse subset of neurons, thereby preventing the presynaptic spectrin signal from being obscured by postsynaptically distributed spectrin. Interestingly, despite the presence of MPS in nearly all axons, the periodic structure was disrupted at most (~70%) of the presynaptic sites imaged (Fig. 4A, B). Next, we imaged βII spectrin distribution in the dendritic spines of cultured hippocampal neurons at DIV 18, using immunofluorescence against endogenous βII spectrin, and observed the periodic structure of βII spectrin in a significant fraction (at least 25%) of the spine necks (Fig. 4C). Notably, the periodic pattern was observed even in some of the spine necks stemming from the shaft regions that exhibited an irregular βII spectrin distribution (Fig. 4C-2).

**Fig. 4.**
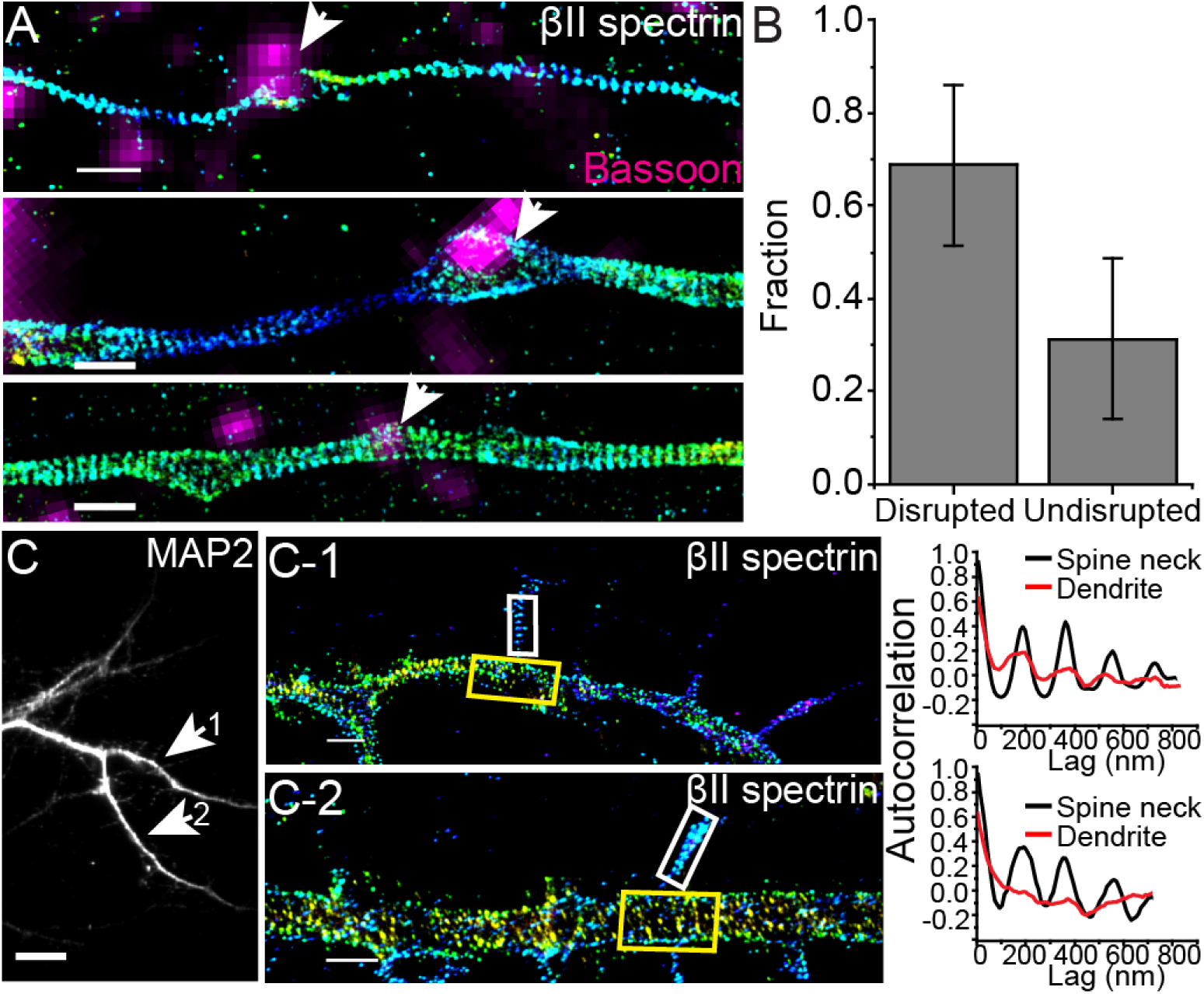
The distribution of βII spectrin in axonal boutons and dendritic spines. (A) Representative STORM images of βII spectrin in axons that contain presynaptic boutons marked by a presynaptic marker, Bassoon (magenta). βII spectrin is labeled by expressing GFP-βII spectrin (or GFP-αII spectrin) in a sparse subset of cultured neurons, and hence some bassoon-positive regions do not overlap with the spectrin-positive axon. Top and middle panels show examples of bassoon-positive presynaptic boutons (arrow indicated), within which the periodic pattern of spectrin is disrupted. Bottom panel shows an example of bassoon-positive presynaptic bouton (arrow indicated), within which the period pattern of spectrin is not disrupted. (B) Bar graph depicting relative fraction of boutons in which the periodic structure is disrupted or undisrupted. (C) The dendritic region of a cultured mouse hippocampal neuron immunostained with MAP2, a dendrite marker. (C-1 and C-2) STORM images of two selected regions showing periodic patterns of immunolabeled endogenous βII spectrin in spine necks, with autocorrelation calculated from the boxed spine neck region (white boxes) and dendritic shaft (yellow boxes). Scale bar: 10 μm for conventional images, 1 μm for STORM images.

### The distribution of βII spectrin in glial processes

We next examined whether the MPS structure also forms in glial-cell processes. Glial cells express βII spectrin, but at a lower level than neurons (35, 36). We cultured glial cells from mouse brain or rat sciatic nerve tissues, used specific immunomarkers to identify four different glial types, including astrocytes, microglia, NG2 glia and Schwann cells, and imaged their endogenous βII spectrin distribution using immunolabeling and STORM. The βII spectrin distribution appeared to be largely irregular in most of the processes of all four glial types imaged, although patches of periodic βII spectrin patterns could be observed in a small fraction of these processes (Fig. 5A and Fig. S7), similar to the appearance of βII spectrin in dendrites. Autocorrelation analysis showed that the degree of periodicity of the βII spectrin distributions in the glial-cell processes was similar among all four glial types, and also similar to that observed in neuronal dendrites, but much smaller than that observed in neuronal axons (Fig. 5B).

**Fig. 5.**
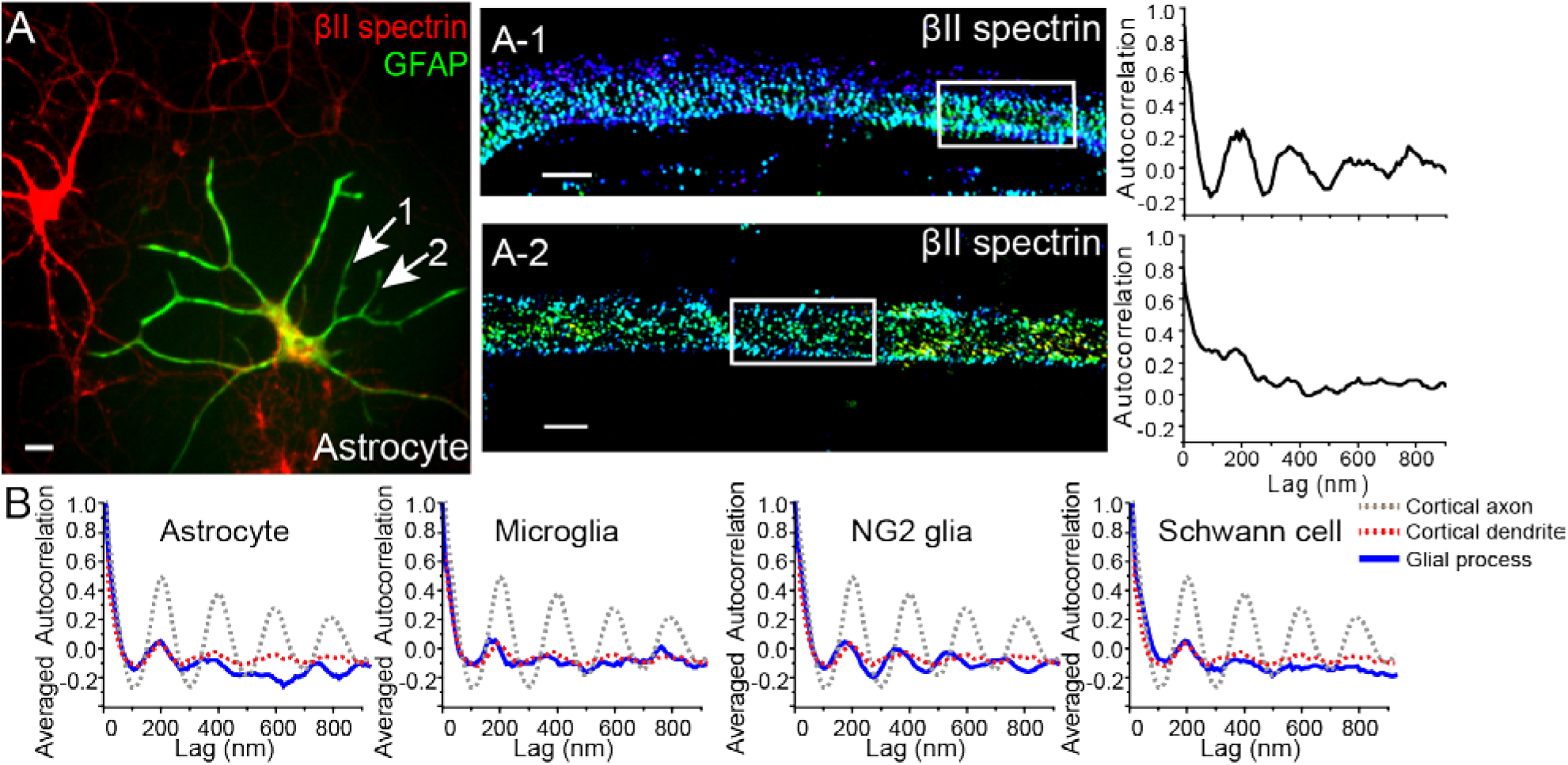
Sparse presence of the periodic βII spectrin structure in glial cell processes. (A) Left: An astrocyte stained for the astrocyte marker GFAP (green) and βII spectrin (red); middle: Two representative STORM images of βII spectrin from two processes of that astrocyte (indicated by arrows) displaying relatively periodic (top) or irregular (bottom) spectrin distribution; right: Autocorrelation analysis of the boxed regions. (B) Averaged autocorrelation functions (blue) calculated from multiple randomly selected processes of astrocytes, microglia cells, NG2 glia, and Schwann cells (N = 10 for astrocytes, N = 9 for microglia, N = 8 for NG2 glia, N = 9 for Schwann cells). For comparison, the dotted gray and red curves show the averaged autocorrelation functions of βII spectrin distribution in axons and dendrites of cortical vGlut1+ neurons, respectively (reproduced from Fig. 1C). Scale bar: 20 μm for conventional image, 1 μm for STORM images.

### The MPS structure was observed in neurons from multiple invertebrate and vertebrate animal species

Spectrin tetramers are evolutionarily conserved with the lengths and major structural domains of the αII and βII spectrin homologs preserved across the metazoan kingdom (Fig. S8) (11, 22). Next, we examined whether MPS can also form in other animal species in addition to rodents. We first studied two additional vertebrate species, *Gallus gallus* (chicken), a non-mammalian vertebrate, and *Homo sapiens.* We examined the endogenous βII spectrin distribution using immunofluorescence labeling and STORM imaging in cultured chicken neurons and human iPS cell-derived motor neurons. βII spectrin exhibited a long-range, periodic pattern along the axonal shafts, with a spacing of ~190 nm in both chicken neurons and human iPS-derived motor neurons (Fig. 6A, B). Next we extended our studies to two invertebrate systems: *C. elegans* and *Drosophila.* To this end, we created fluorescent fusion proteins of β spectrin for *C elegans* (UNC-70) and *Drosophila* (dβ-Spectrin) and over-expressed these fusion constructs with neuronal-specific promotors in *C elegans* and in glutamatergic neurons of *Drosophila* larva (see Supporting Information). We used structured-illumination microscopy (SIM) (37) to image these neurons *in vivo*, either in the whole animal (*C. elegans*) or in brain tissues (*Drosophila*), rather than in cultured neurons. In both *C. elegans* and *Drosophila* neurons, the expressed β spectrin proteins adopted a periodic structure with a spacing of ~190 – 200 nm (Fig. 6C, D), consistent with the conserved lengths and domain structures of spectrin tetramers across different species. These results indicate that the MPS structure can form not only in vertebrate neurons but also in invertebrate neurons. However, we note that, although over-expression of βII spectrin does not appear to perturb the formation of the MPS structure in rodent axons, such over-expression does promote MPS formation in dendrites of rodent neurons (16). Therefore, future experiments on imaging endogenous spectrin are needed to confirm whether the MPS structure is formed by endogenous proteins in *C. elegans* and *Drosophila.* Because spectrin is expressed in many neuronal and non-neuronal cells, the resulting high background makes such imaging experiments difficult using immunofluorescence. Using genome editing to introduce fluorescent tags to the endogenous spectrin in specific populations of neurons provides a promising solution.

**Fig. 6.**
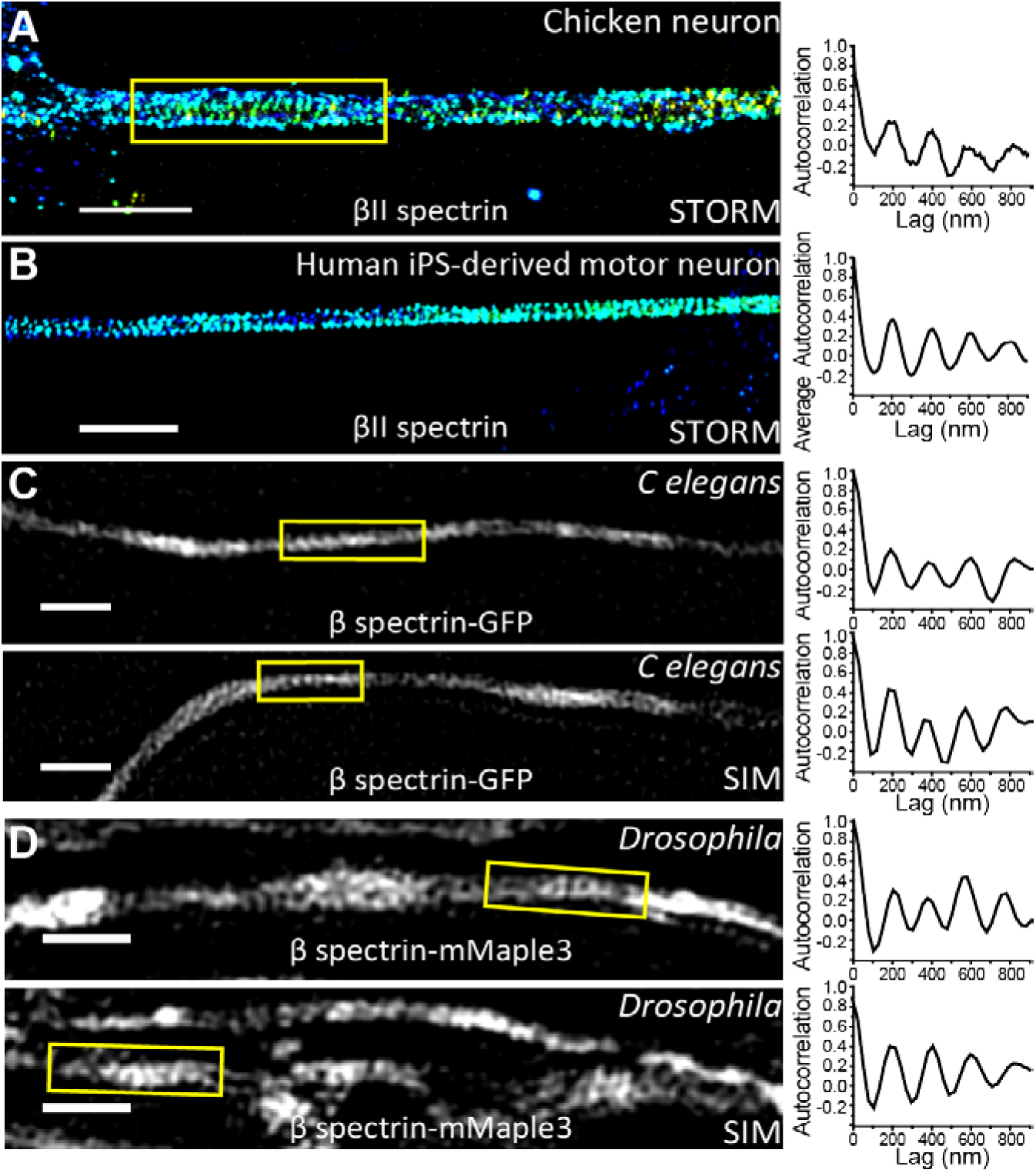
The observation of MPS in neurons across different animal species. (A) Representative STORM image of immunolabeled βII spectrin in the axon of a cultured chicken neuron (left) and the autocorrelation function calculated from the boxed region of this image (right). Scale bar: 2 μm. (B) Representative STORM image of immunolabeled βII spectrin in the axon of a human iPS cell-derived motor neuron (left) and the autocorrelation function calculated from this image (right). Scale bar: 2 μm. (C) Two representative SIM images of β spectrin (UNC-70)-GFP in *C. elegans* neurons imaged directly in the animals (left), and the corresponding autocorrelation function of the boxed regions (right). Scale bar: 1 μm. (D) Two representative SIM images of β spectrin-mMaple3 in *Drosophila* neurons imaged in *Drosophila* brain tissue (left), and the corresponding autocorrelation function of the boxed region (right). Scale bar: 1 μm.

## Discussion

We have recently discovered that actin, spectrin and associated molecules form a membrane-associated periodic skeletal structure in the axons of rodent neurons (12). Here, we show that this actin-spectrin-based MPS structure is prevalently present in a wide spectrum of neuronal cell types cultured from the rodent central and peripheral nervous systems, including excitatory and inhibitory neurons in cortex, hippocampus, and midbrain, and many specific neuronal subtypes, such as granule cells, pontine nuclei neurons, dopaminergic neurons, olfactory neurons, Parvalbumin cells, Golgi cells, Purkinje cells, motor neurons and sensory neurons. While we were finalizing our paper, an independent study reported the presence of periodic actin structure in several neuronal types in the rodent central and peripheral nervous systems (23), and the list of cell types reported in (23) partially overlap with the neuronal types investigated here. Using specific neuronal markers to distinguish neuronal cell types, we further showed that in all these neuronal types, the MPS structure preferentially forms in axons compared to dendrites. Interestingly, despite the prevalent presence of MPS in nearly all axons, we showed that this periodic structure is disrupted in most presynaptic boutons. This observation suggests the possibility that MPS may inhibit synapse formation and thus needs to be disrupted for bouton formation, and that the small fraction of boutons that maintained the periodic structure might be relatively immature synapses. On the other hand, the periodic structure is formed in a significant fraction of dendritic spine necks, including ones that stemmed from the dendritic shafts lacking the MPS structure, suggesting a specific mechanism to promote the formation of this periodic structure in the spine neck. We also examined four types of rodent glial cells, including astrocytes, microglia, NG2 glia and Schwann cells and observed isolated patches of periodic spectrin distributions in a small fraction of glial-cell processes, consistent with the observation of periodic actin and spectrin distributions in rodent oligodendrocytes (23). It is interesting to note that this periodic structure appeared to be absent in the majority of glial-cell processes in our images, and our quantitative analysis showed that the degree of periodicity for spectrin distributions in glial processes was similar to that observed in neuronal dendrites and much smaller than that observed in neuronal axons. Because the expression level of βII spectrin in glial-cell processes is similar to that in dendrites and lower than that in axons (35, 36), this observation is consistent with the notion that local concentrations of βII spectrin is a key determinant for the MPS formation (16). Taken together, these results suggest that the MPS structure might have some tendency to self-assemble, but the axon is a specialized cellular compartment where the formation of MPS is strongly promoted. Whether this substantially higher propensity of MPS formation in neuronal axons observed in all examined cultured nervous system cell types can be extended to *in vivo* systems awaits future investigations. Finally, we found that spectrin is capable of adopting a periodic distribution in neurons from a wide variety of animal species across both invertebrates and vertebrates, ranging from *C. elegans* to *H. sapiens*, suggesting that the MPS structure may be an evolutionally conserved structure in neurons across different species. The high prevalence of this structure in the axons of diverse neuronal types and animal species suggests that the actin-spectrin-based MPS is a key functional component of the axon. Indeed, such a periodic skeletal structure formed by actin rings connected by flexible spectrin tetramers provides a robust and flexible support for the axonal membrane, which could be particularly important for maintaining the integrity of axons under mechanical stress and can explain why spectrin depletion results in axon buckling and breakage during animal locomotion (4, 6). The ability of this periodic sub-membrane skeleton to organize membrane proteins, such as ion channels (12), into a periodic distribution might also impact the generation and/or propagation of action potentials as well as other signaling pathways at the axonal membrane.

### Materials and Methods

Procedures for cell culture, antibodies and immunofluorescence are described in Supporting Information (SI). STORM imaging was performed as previously described (12, 14). Briefly, labeled cells were imaged with continuous illumination of 657-nm laser for imaging and 405 nm for activation of the photoswitchable dyes in an oblique-incidence geometry on an Olympus IX-71 inverted microscope. STORM images were generated from a sequence of about 20,000~40,000 image frames at a frame rate of ~60 Hz. SIM imaging (37) was performed on a Zeiss Elyra super-resolution microscope in the Center for Biological Imaging at Harvard University. SIM imaging analyses were performed with the provided software for the Zeiss Elyra system. Autocorrelation analyses were performed as previously described (12, 16). The autocorrelation amplitude is defined as the difference between the first peak (at ~ 190 nm) and the average of the two first valleys of the autocorrelation curve. See SI Materials and Methods for more details.

## Acknowledgements

This work is supported in part by Howard Hughes Medical Institute (to X.Z. K.S. and M.F.), the National Institutes of Health (NS089786 to M.T.L, NS048392 to K.S. and NS053538 to M.F.), and the ALS Association (to T.M.). R.Z. is an HHMI Fellow of the Life Sciences Research Foundation. Z.W. is supported in part by a Bristol-Myers Squibb Postdoctoral Fellowship at the Rockefeller University. X.Z., K.S., M.F. are Howard Hughes Medical Institute investigators.

